# Time-course single-cell RNA sequencing reveals transcriptional dynamics and heterogeneity of limbal stem cells derived from human pluripotent stem cells

**DOI:** 10.1101/2020.09.29.319756

**Authors:** Changbin Sun, Hailun Wang, Qiwang Ma, Chao Chen, Jianhui Yue, Bo Li, Xi Zhang

**Affiliations:** BGI-Shenzhen, Shenzhen, 518083, China; BGI Education Center, University of Chinese Academy of Sciences, Shenzhen 518083, China; China National GeneBank, BGI-Shenzhen, Shenzhen 518082, China; Department of Radiation Oncology, School of Medicine, Johns Hopkins University, Baltimore, MD 21218, United States; Section of Cell Biology and Physiology, Department of Biology, University of Copenhagen, Copenhagen, Denmark

**Keywords:** LSCs, scRNA-seq, identity, developmental trajectory

## Abstract

Human pluripotent stem cell-derived limbal stem cells (hPSC-derived LSCs) provide a promising cell source for corneal transplants and ocular surface reconstruction. Although recent efforts in the identification of LSC markers have increased our understanding of the biology of LSCs, the lack of knowledge of the developmental origin, cell fate determination, and identity of human LSCs hindered the establishment of differentiation protocols for hPSC-derived LSCs and hold back their clinical application. Here, we performed a time-course single-cell RNA-seq to investigate transcriptional heterogeneity and expression changes of LSCs derived from human embryonic stem cells. Based on current protocol, expression heterogeneity of reported LSC markers were identified in subpopulations of differentiated cells. EMT has been shown to occur during differentiation process, which could possibly result in generation of untargeted cells. Pseudotime trajectory analysis revealed transcriptional changes and signatures of commitment of hPSCs-derived LSCs and their progeny - the transit amplifying cells. Furthermore, several new makers of LSCs were identified, which could facilitate elucidating the identity and developmental origin of human LSCs *in vivo*.

## Introduction

Human limbal stem cells (LSCs) are located in a narrow area around the cornea and connect directly to the sclera (Busin et al., 2016; Davanger and Evensen, 1971; Kawakita et al., 2011). Other than self-renewal capability for homeostasis maintenance, LSCs have unipotency to differentiated into cornea epithelial cells and play vital role in corneal regeneration and repair to sustain the corneal integrity and homeostasis (Ebrahimi et al., 2009). However, internal or external factors, such as genetic, chemical, burn, infection etc., could result in limbal malfunction and limbal stem cells deficiency (LSCD), and lead to reduced vision and blindness (Barut Selver et al., 2017; Kim and Mian, 2017; Le et al., 2018).

Among different treatment options, LSC transplantation is currently the best curative treatment that can improve both vision and quality-of-life in patients with ocular surface disorders caused by LSCD (Atallah et al., 2016). Although the developmental origin of LSC remains enigmatic, most studies considered that the corneal epithelium descend from surface ectoderm (SE) (Gonzalez et al., 2018; Hongisto et al., 2017), and give rise to structures of the epidermis and ectodermal associated appendages such as hair, eye, ear, and the mammary gland etc. (Tchieu et al., 2017). However, developmental surface ectodermal cells and their derivatives are difficult to isolate and study in human. Our understanding on cell-fate specification of the limbal stem cells *in vivo* are limited and largely from studies of classic model organisms, such as mouse (Kaplan et al., 2019; Wolosin et al., 2004) and *Xenopus* frogs (Sonam et al., 2019). But it is well-known that final maturation pathways are significantly different between humans and other model animals, though their pre-implantation development appears relatively similar (Rossant, 2015). Thus, the directed differentiation of human pluripotent stem cells (hPSCs) to LSCs could offer an alternative model system to explore these cells’ identity and fate decisions for basic and clinical applications (Ahmad et al., 2007; Chakrabarty et al., 2018; Hanson et al., 2013; Hayashi et al., 2016; Kamarudin et al., 2018; Tchieu et al., 2017). However, available differentiation protocols are still inefficient and suffer from excessive heterogeneity (Pattison et al., 2018). The lack of specific markers for LSCs, and our limited knowledge about intrinsic signaling cascades and developmental mechanisms of human LSCs hindered the clinical application of LSCs (Chakrabarty et al., 2018; Gonzalez et al., 2018).

Single-cell RNA sequencing (scRNA-seq) is a powerful tool to quantify transcripts in individual cells to understand gene expression changes at single-cell resolution (Gurtner et al., 2018). Since the first publication in 2009 (Tang et al., 2009), scRNA-seq have increasingly been utilized in many fields, such as developmental biology to delineate cell lineage relationships and developmental trajectories (Clark et al., 2019; Hu et al., 2019; Su et al., 2017). In this study, we performed a time-course single-cell transcriptomic analysis of LSCs derived from human embryonic stem cells *in vitro* to understand transcriptional regulation during human LSCs development.

## Experimental Procedures

### Cell culture

The Ethics Committee of BGI-IRB approved this study. Human ESC lines H9 were cultured as previous description (Sun et al., 2018). Briefly, cells were retrieved from liquid nitrogen tank and cultured in hESC medium (DMEM/F12 basic medium (Life Technologies), 20% knockout serum replacement (KSR, Life Technologies), 1×L-glutamine (Life Technologies), 1×MEM NEAA (Life Technologies), 0.1mM 2-Mercaptoethanol (Life Technologies) and 50 ng/mL human FGF-2 (Life Technologies)) on mitomycin C (Sigma) treated murine embryonic fibroblasts (MEFs). To sustain undifferentiated states, cells were fed daily with fresh medium. For passaging, colonies were dispersed into small clumps with 1mg/mL Collagenase IV (Life Technologies) for 20 min at 37℃, then plated onto Matrigel hESC-qualified Matrix (Corning)-coated dishes in mTeSR1 medium (Stemcell Technologies) at a ratio of 1:3 to 1:6. In the feeder-free medium, ReLeSR^TM^ (Stemcell Technologies) were used for dissociation and passaging according to the manual.

### LSCs induction

LSCs were differentiated from human ESCs according to the published protocols with some changes (Hongisto et al., 2017; Mikhailova et al., 2014). Briefly, when colonies reaching about 80-90% confluency, ReLeSR^TM^ were used to digest cells into clumps. Then, these clumps were suspended in LSCs induction medium (DMEM/F12 basic medium, supplemented with 20% KSR, 1×L-glutamine, 1×MEM NEAA, 0.1mM 2-Mercaptoethanol) adding 10μM Y-27632 (Sigma) at 37℃ to induce embryoid body (EB) formation overnight. For LSCs differentiation, EBs were cultured in LSCs induction medium supplemented with 10 μM SB-505124 and 50 ng/ml FGF-2 for 1 day. Then, medium changed with LSCs induction medium supplemented 25 ng/ml bone morphogenetic protein 4 (BMP4) (R&D) for 2 days. Thereafter, the induced cultures were seeded onto plates coated with 0.75 μg/cm2 LN521 (BIOLAMINA) and 5 μg/cm2 col IV (Sigma) in a defined and serum-free medium CnT-30 (CELLNTEC). For next days before collection for scRNA-seq, the cells were maintained in CnT-30 and change the medium every 3 days.

### scRNA-seq library construction and sequencing

scRNA-seq experiments were performed by Chromium Single Cell 5’ Library & Gel Bead Kit (10x Genomics), according to the manufacturer’s protocol. Briefly, cells were digested with TrypLE™ Select (ThermoFisher Scientific) and single cell suspension were harvested, washed with PBS twice, and filtered by 40 μm cell strainers (BD Falcon) before Gel Bead-In Emulsions (GEMs) generation and barcoding. Single-cell RNA-seq libraries were obtained following the 10x Genomics recommended protocol, using the reagents included in the kit. Libraries were sequenced on the BGISEQ-500 (BGI) instrument (Natarajan et al., 2019) using 26 cycles (cell barcode and UMI (Islam et al., 2014)) for read1 and 108 cycles (sample index and transcript 5’ end) for read2.

### scRNA-seq Analysis

#### Quality control

The scRNA-seq data was processed using cellranger-3.0.2 (https://support.10xgenomics.com) for each sample with default parameters mapping to the human GRCh38 genome expect the number of recovered cells (--expect-cells option) was set to 8 000.

For each library, we filtered outlier cells using the median absolute deviation from the median total library size (logarithmic scale), total gene numbers (logarithmic scale), as well as mitochondrial percentage, as implemented in scran, using a cutoff of 3 (isOutlier, nmads = 3) (Lun et al., 2016). For filtering lowly or none expressed genes, genes expressed across all the cells detected in less than 10 cells were removed, and totally 22 501 genes were kept for downstream analysis. Then, clean gene-cell UMI count matrix was loaded as Seurat object using R package Seurat 3.0 (Macosko et al., 2015) or cds object using R package monocle 3 (Cao et al., 2019) to manage our dataset for the further analysis with default parameters otherwise will be mentioned in detail.

#### Cell cycle phase assignment

To assign cell cycle phase for each cell, cell cycle scores (i.e., G2/M scores and S scores) and phases (i.e. G1, G2/M, and S) for each cell on the basis of scores using function CellCycleScoring from R package Seurat based on the expression levels of a panel of phase-specific marker genes (Nestorowa and Hamey, 2016).

#### Normalization and dimension reduction

The quality control dataset were then analyzed using the Seurat v.3.0 pipeline with NormalizeData function to normalize our data, FindVariableFeatures funtion to assign top 2000 highly variably expressed genes, ScaleData function of argument vars.to.regress to remove confounding sources of variation (variables to regress out including mitochondrial mapping percentage, number of UMI). Following normalization and scaling, RunPCA function were performed to capture principal components using the top 2000 highly variably expressed genes. UMAP was applied to visualize and explore data in two-dimensional coordinates, generated by RunUMAP function in Seurat.

#### Cell cluster

For cell clustering, a graph-based clustering approach (Macosko et al., 2015) were used to cluster the cells into candidate subpopulations. The first 50 PCs in the data were applied to construct an SNN matrix using the FindNeighbors function in Seurat v3 with k.param set to 20. We then identified clusters using the FindClusters command with the resolution parameter set to 0.5.

#### Differential Expression Analysis

To find differential expressed genes (DEGs), Wilcoxon Rank Sum test were performed for significant test using Seurat function FindAllMarkers for every cluster compared to all remaining cells and FindMarkers for distinguishing each other. Genes with average natural log fold change more than 0.25 and FDR less than 0.01 were assigned as DEGs.

#### Pseudotime trajectories analysis

For pseudotime trajectories analysis, the quality control dataset with cell clustering information were analyzed using the monocle3 (http://cole-trapnell-lab.github.io/monocle3/) pipeline. The new_cell_data_set function in the package was used to create cds object, and preprocess_cds function was applied for data normalization and principal component analysis with num_dim setting to 50. Then, reduce_dimension, cluster_cells, and learn_graph functions were used for data reduction, cell clustering, and pseudotime trajectories construction, respectively. UMAP was applied to visualize and explore data in two-dimensional coordinates using plot_cells function.

## Results

### Single-cell RNA sequencing revealed expression heterogeneity of hESC-derived LSCs

Human embryonic stem cell (H9) was used to differentiate to LSCs via a surface ectodermal stage (Hongisto et al., 2017; Mikhailova et al., 2014) (Fig S1A). To characterize obtained hESC-derived LSCs, we performed scRNA-seq at four time points, Day 0 before induction, Day 7, Day 14, and Day 21 after induction. In total, 18541 cells were sequenced, and data from 14241 cells were used for the following analysis after filtering out low quality cells, including 4687 cells, 4784 cells, 3210 cells, and 1560 cells from Day 0, Day 7, Day 14, and Day 21, respectively (Fig S1B-S1E).

Gene expression analysis showed that, at Day 0, POU5F1, SOX2, and NANOG were highly expressed in most cells, accounting for 99.98 %, 99.73 %, and 82.27% of all the analyzed cells, respectively (Fig 1A and 1E), which indicated that these cells used for hESC-derived LSCs differentiation were pluripotent.

**Fig 1.**
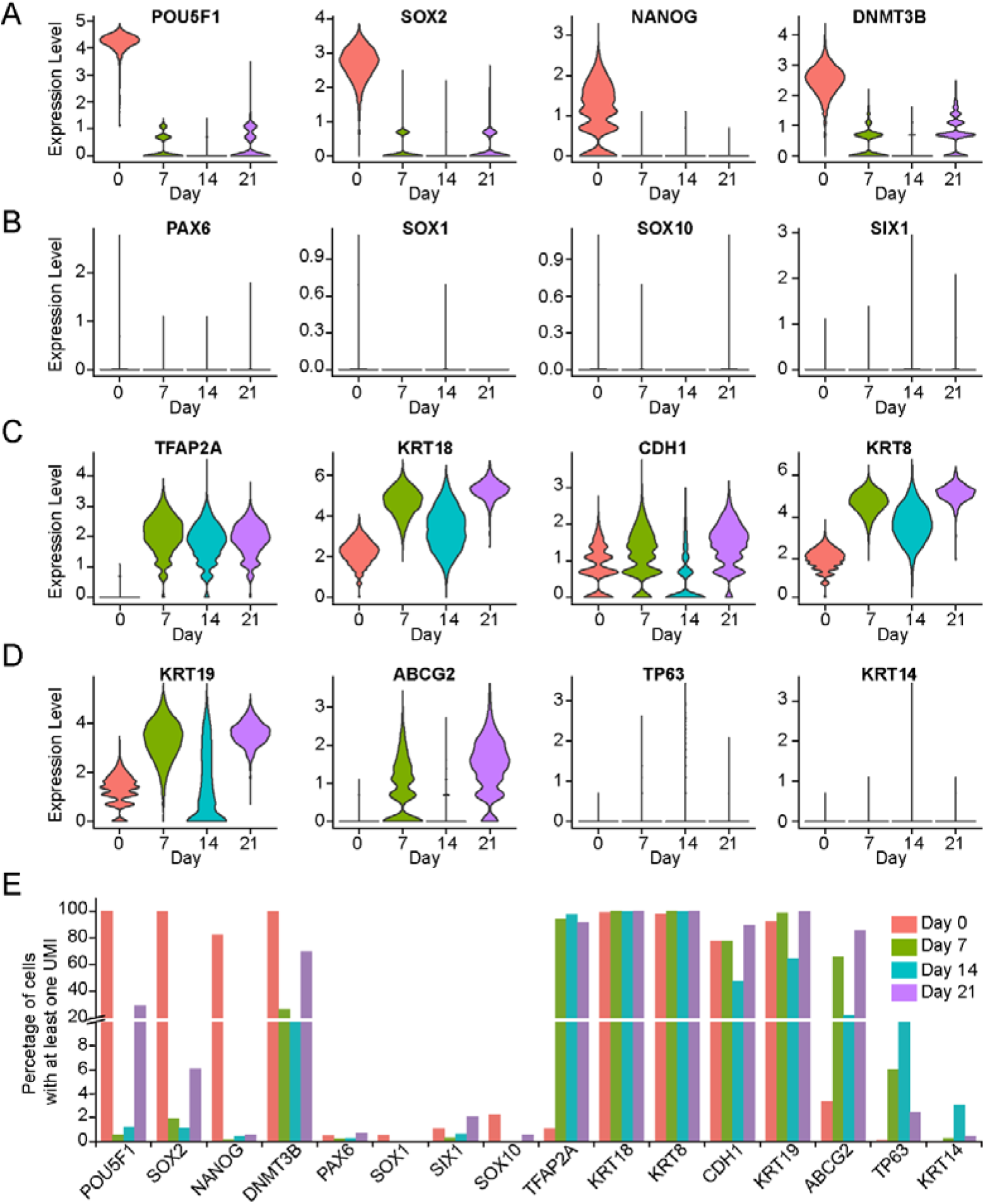
Single-cell RNA sequencing analysis of human embryonic stem cells-derived LSCs differentiation at different time point. (A-D) Violin plot representing expression of pluripotency (A), neural ectoderm (B), surface ectoderm and epithelium (C), and candidate LSCs (D) markers at the four times. (E) Barplot representing percentage of cells expressed the selected pluripotency, neural ectoderm, surface ectoderm and epithelium, and candidate LSCs markers at the four times.

At Day 7, 94.25% of cells expressed transcription factor TFAP2A while only a few of the cells expressed pluripotency markers (POU5F1 (0.57%), SOX2 (1.94%), and NANOG (0.14%)) neuroectodermal markers (SOX1 (0.00%) and PAX6 (0.24%)), neural crest marker SOX10 (0.04%), and cranial placode marker SIX1 (0.32%) (Fig 1A, 1B and 1E), demonstrating the residual pluripotency and a direction of differentiation toward surface ectodermal progenitors (Tchieu et al., 2017). In addition, a range of both epithelial progenitor and candidate LSCs markers (Gonzalez et al., 2018), such as KRT19, KRT18, TP63 (p63), CDH1, and ABCG2, were expressed in this population. However, some of these genes showed high expression variability between clusters (Fig 1B, 1C and 1E). For example, TP63 (well-known as p63), which has been linked to successful limbal transplantation (Rama et al., 2010), expressed in a small portion of cells (6.034%) (Fig 1E).

At Day 14 and Day 21, we found the expression of epithelial progenitor and candidate LSCs markers were highly variable as well. Percentage of cells expressing TP63 decreased from 11.21% at Day 14 to 2.46% at Day 21 (Fig 1E). In contrast, most cells (85.67%) at Day 21 expressed ABCG2, one of the widely used makers of LSCs (Budak et al., 2005; de Paiva et al., 2005; Gonzalez et al., 2018; Vattulainen et al., 2019), while only 21.82% of cells at Day 14 had ABCG2 expression. Furthermore, several markers of terminally differentiated LSCs, such as KRT3 and KRT12, were not detected in any cells at Day 14 and Day 21, indicating that these cells were still at immature differentiation stages.

### Time-course Single-cell RNA-seq profiling showed specific changes of gene expression during hESCs-LSCs differentiation

To investigate transcriptional changes during hESCs-LSCs differentiation, we integrated data from the four time points for dimension reduction and visualization. Results showed that all cells were grouped into 11 clusters (Figure 2A). Among the clusters, cluster 2 and 3 are all from Day 0 (Figure 2B). Not surprisingly, these cells exhibited highest expression of pluripotent genes POU5F1, SOX2, NANOG, and DNMT3B (Fig 2D). In contrast, expression of surface ectodermal genes, such as TFAP2A, TFAP2B, TFAP2C, HAND1, GATA3, IFR6, WISP1, and NR2F2, were upregulated throughout differentiation (Figures 2D). Unexpectedly, epithelial genes such as CDH1, EPCAM, KRT8, and KRT18, were lowly expressed in cluster 1, while mesenchymal genes such as CDH2, COL1A1, COL1A2, and FBN1 were highly expressed, indicating that cluster 1 were mesenchymal cells. In addition, neural genes such as COL2A1, SOX11, OTX1 and SIX1 were upregulated in cluster 9. Also, cells from Day 0 and Day 21 were separated, whereas some cells from Day 7 and Day 14 were clustered with each other (Figure 2A and 2B), indicating these cells at Day 7 and Day 14 had similar expression profiles. Therefore, these results demonstrated that during hESCs to LSCs differentiation, hESCs gave rise to cells with none epithelial characteristics.

**Figure 2.**
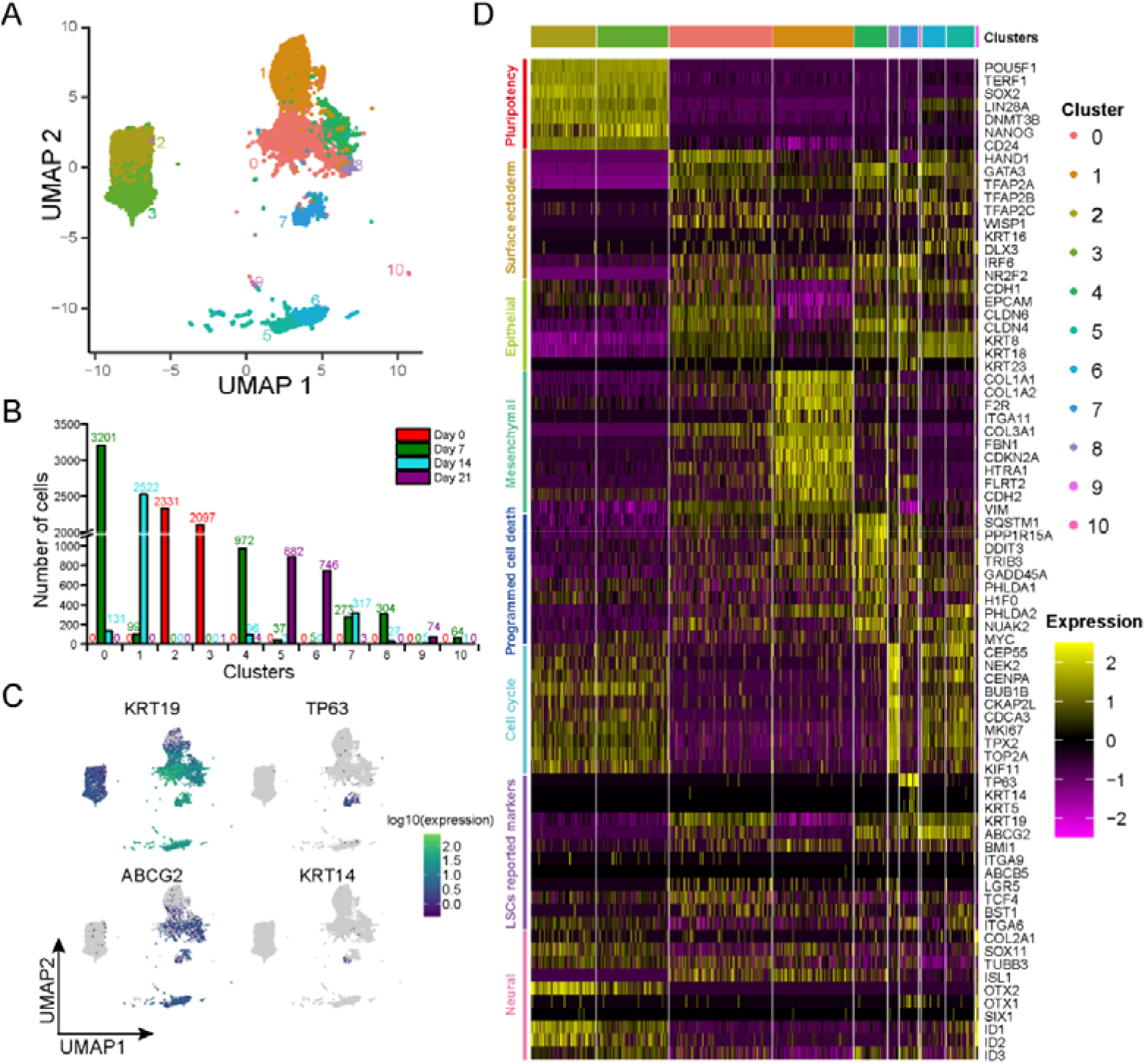
Time-course Single-cell RNA sequencing profiling reveals heterogeneity by current hESCs-derived LSCs differentiation method. (A) UMAP visualizing the results of clustering for cells sequenced at the four times. (B) Barplot showing number of cells in the four days for each cluster. (C) Feature plots visualizing the four key LSCs marker genes expression in the cells. (D) Heatmap representing genes differentially expressed among the clusters.

Notably, differential gene expression (DGE) analysis showed that genes related to cell cycle and programmed cell death were highly expressed in cluster 8 and cluster 4, respectively (Fig 2D). In cluster 8, expression of genes such as TOP2A, MKI67, TPX2, BUB1B, and CEP55 were significantly upregulated, while SQSTM1, DDIT3, PPP1R15A, H1F0, and TRIB3 etc. showed higher expression in cluster 4 (Fig 2D). To avoid the potential bias from cell cycle effects, we assigned cell cycle phase to each cell. Then, we only extract cells in clusters with G2M phase to compared cycle related genes expression. Results demonstrated that cycle related genes, such as TOP2A, MKI67, TPX2, BUB1B, and CEP55 ect., were highly expressed in cluster 8 as well (Fig S2D and S2E). These results demonstrated that no obvious cell cycle effects on data dimension reduction, and cell cycle effects did not obviously impact cell clustering, and cluster 8 are indeed highly expanding cells.

Next, we investigated expression of several putative LSC-associated markers (e.g. KRT19, ABCG2, VIM, ITGA9, TP63, KRT14, KRT15, KRT5) and differentiation-associated markers (e.g. KRT3 and KRT12) (Gonzalez et al., 2018; Schlotzer-Schrehardt and Kruse, 2005) during hESC-LSCs differentiation. Results shown that differentiation-associated markers KRT3 and KRT12 were not detected in all clusters. Interestingly, putative LSC-associated markers TP63 and KRT14 were highly expressed in cluster 7 while KRT19 and ABCG2 were upregulated in all the clusters except cluster 1 (Fig 2C and 2D). Taken together, these results indicated that cells in cluster 0, cluster 5, cluster 6, cluster 7, cluster 8, and cluster 10 could be progenitors of LSCs, LSCs and their progeny in the different stages of development.

### Pseudotime analysis revealed unique hESC-LSCs developmental trajectory

To investigate hESC-LSCs developmental trajectory, we performed pseudotime analysis to study the path and progress of individual cells undergoing hESCs-derived LSCs differentiation (Trapnell et al., 2014). The resultant trajectory indicated that a trifurcation point in cluster 0 could lead to cells fate commitment toward cluster 1 (Branch 1), cluster 4 (Branch 2), and cluster 8 that further differentiate to cells in cluster 7, 10, 5 and 6 (Branch 3) (Fig 3A and 3B).

**Figure 3.**
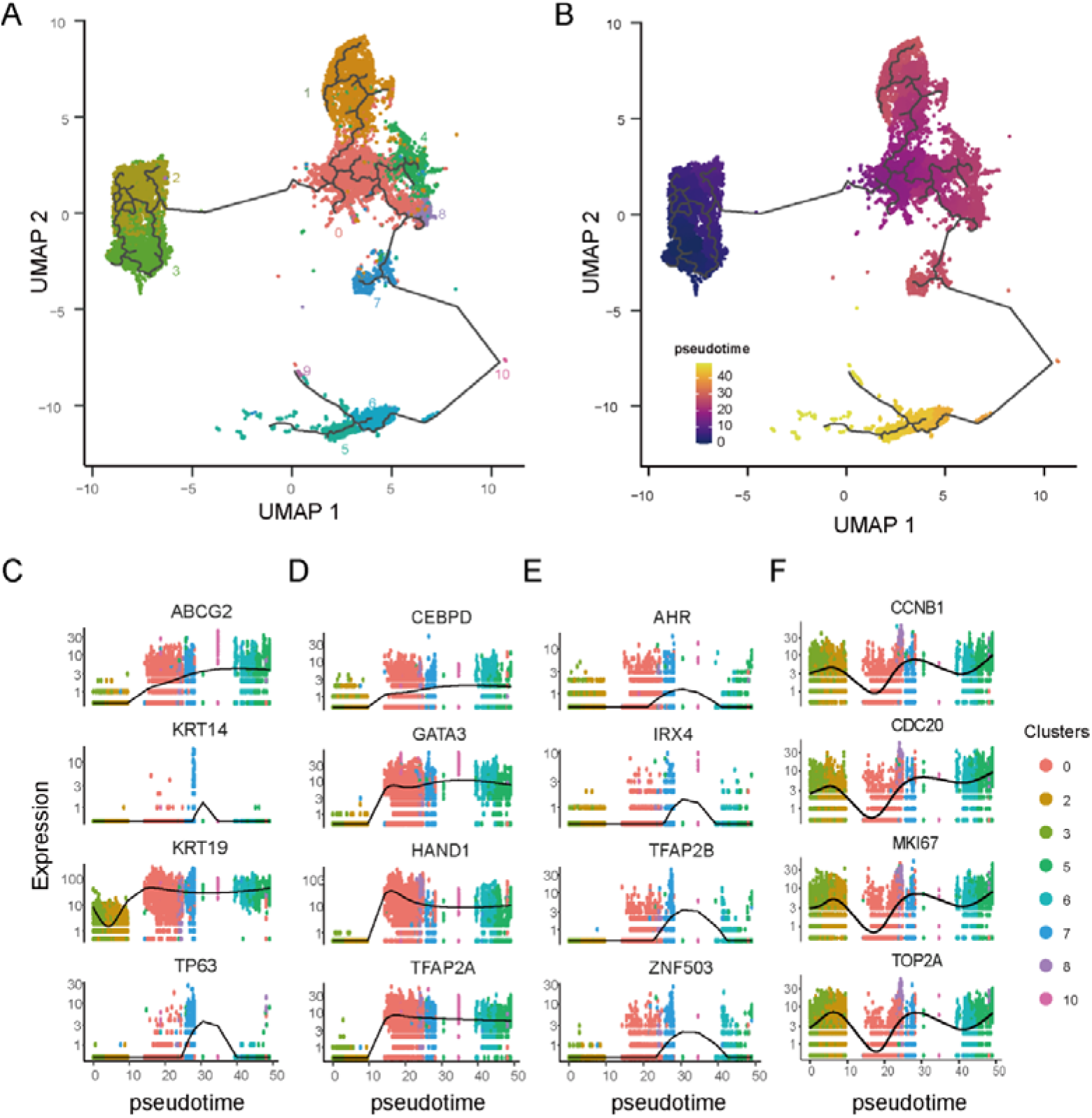
Pseudotime analysis characterizes expression changes throughout hESCs-derived LSCs differentiation. (A and B) UMAP visualizing developmental trajectories of cells in each cluster (A) and pseudotime assigned to each cell. (C) Plotting showing tracking changes of the four LSCs marker genes over hESC-derived LSCs differentiation pseudotime. (D and E) Plotting representing tracking changes of TFs upregulated upon differentiation and continually highly expressed (D), and upregulated in certain period (E) over hESC-derived LSCs pseudotime. (F) Plotting representing tracking changes of cell cycle related genes over hESC-derived LSCs pseudotime.

In Branch 1, CDH1 (E-cadherin) and CDH2 (N-cadherin), two well-known cadherins, were differently expressed between cells in cluster 0 and cluster 1 (Fig S3B and S3C). Specifically, CDH1 was upregulated in cluster 0 while CDH2 was expressed significantly higher in cluster 1. The loss of epithelial surface marker CDH1 and the acquisition of mesenchymal marker CDH2 is considered as the hallmark of epithelial-mesenchymal transition (EMT), which play pivotal role in developmental regulation, such as Neural Crest Formation (Kim et al., 2017). Additionally, upregulated genes in cluster 1 compared to cluster 0 were overrepresented significantly in nervous system development (Fig S3D), indicating the possible generation of neural crest like cells during the hESCs to LSCs differentiation process. In Branch 2, cells were undergoing programmed cells death (apoptosis) as mentioned in above section (Figure S3D). Apoptosis is a positive regulator of stem cells populations, it plays fundamental roles in development and tissue homeostasis (Fuchs and Steller, 2011; Kaplan et al., 2019). Branch 3 identified the main hESC-LSCs developmental trajectory. Epithelium development and epithelial cell proliferation related genes were upregulated in the Branch 3 differentiation process (Figure S3D). Increasing expression of candidate LSC markers KRT19, ABCG2, KRT14, and TP63 were seen in cluster 5, cluster 6, and cluster 7 (Figure 2C). Pseudotime analysis further demonstrated that these candidate markers exhibiting different trajectory patterns in Branch 3 (Figure 3C). In addition, some transcription factors (TFs), such as CEBPD, GATA3, HAND1, and TFAP2A, were upregulated upon differentiation and stably expressed at high level (Figure 3D), while some TFs, such as AHR, IRX4, TFAP2B, and ZNF530, only upregulated in a certain period of time like TP63 (Figure 3C,E), indicating their distinct roles in hESC-LSCs development. Interestingly, cell cycle related genes, such as CCNB1, CDC20, MKI67, and TOP2A, shown regular oscillations patterns across hESC-LSCs developmental pseudotime to regulate the cell proliferation (Fig 3F).

### Transcriptional difference of subpopulations in hESCs-derived LSCs

In the pseudotime analysis, cluster 7 expressed most reported candidate LSC markers, including TP63 (Pellegrini et al., 2001), KRT14 (Kurpakus et al., 1994), KRT15 (Yoshida et al., 2006), ITGA6 (Hayashi et al., 2008) etc. (Figure S4A). To investigate expression differences among subpopulations in hESCs-derived LSCs, we performed two-two comparisons among cluster 5, cluster 6, and cluster 7 (Figure S4B). Differential expression analysis demonstrated that upregulated genes in cluster 5 and cluster 6 population shown significant enrichment in cell cycle process. In addition, genes involved in cell migration regulation were highly expressed in cluster 5 compared to cluster 6, including cadherin genes CDH5 and CDH13, Integrin genes ITGA2, ITGA6, ITGA3, ITGB6, ITGB1, ITGA5 and ITGAV, collagen genes COL4A1, COL4A2, COL1A2 and COL3A1, and transcription factors SOX9, MYC, STAT3 ect. “X, Y, Z hypothesis” of corneal epithelial maintenance suggested that proliferation of basal cells (X) and migration of centripetal cells replace cells’ lost from the ocular surface (Z) to support the corneal epithelial homeostasis (Thoft and Friend, 1983), indicating cellular and functional variables for corneal epithetical balance. Within the cornea, nuclear p63 (TP63) is expressed by the basal cells of the limbal epithelium, but not by TA cells covering the corneal surface (Pellegrini et al., 2001). Therefore, these results suggested that cells in cluster 7 (TP63 expression) give rise to cells (TACs) in cluster 5 and cluster 6, both of which are the progeny of LSCs exhibiting high, but limited proliferative activity (Beebe and Masters, 1996; Pellegrini et al., 2001).

To identify potential markers to distinguish these cells, we focused on transcription factors (TFs) and cluster of differentiation (CD) genes differentially expressed in cells in cluster 5, cluster 6, and cluster 7. Among the TFs that play key roles in cell fate decision, CXXC5, IRF6, SKIL, RUNX1 etc. as well as TP63 were upregulated in cluster 7. GATA3, EPAS1, HAND1, HOXB2, and CEBPD ect. were highly expressed in cluster 6, while NFE2L3, EVT4, YBX1, FOSL1, and MYC ect. were enriched in cluster 5 (Fig 4D). As to the CD genes, SDC1, ITGB4, CD9, IGF1R, JAG1, CD46, CD151 ect. were highly expressed in cluster 7, while LIFR, CD99, FGFR2, ABCG2 etc. were upregulated in cluster 6, and ENPEP, THY1, CD40, CD44, CDH5 etc. exhibited highest expression in cluster 5 (Fig 4E). All these candidate markers identified here would be valuable for future characterization of different cell types in human cornea.

**Figure 4.**
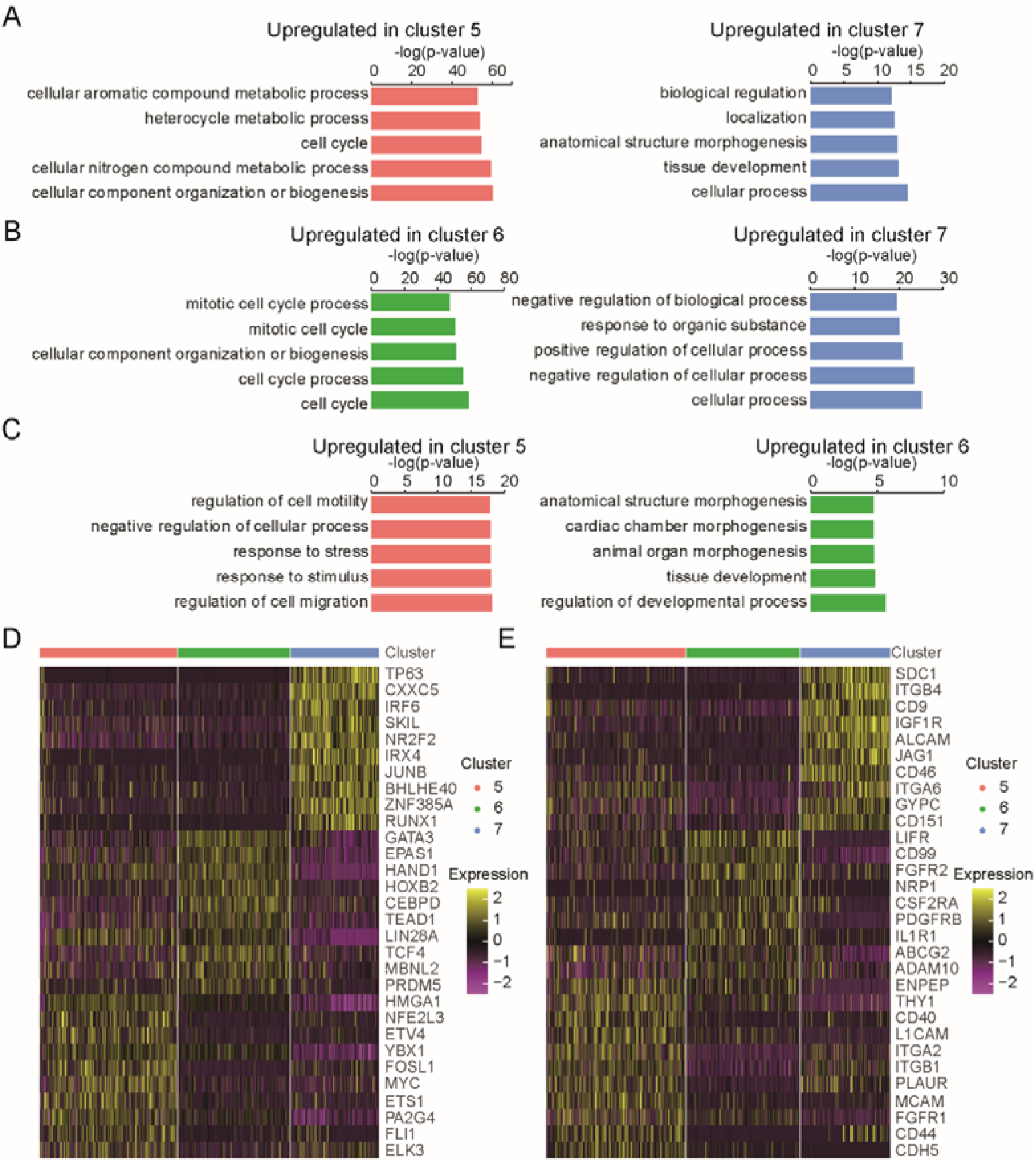
Transcriptional difference of subpopulations in hESCs-derived LSCs. (A-C) Barplots showing GO biological process enrichment for upregulated genes compared between cluster 5 and cluster 7 (A), between cluster 6 and cluster 7 (B), and between cluster 5 and cluster 6 (C). Five terms with lowest p-value were presented. (D) Heatmap representing differentially expressed TFs among cluster 5, cluster 6, and cluster 7. (E) Heatmap representing differentially expressed top CD genes among cluster 5, cluster 6, and cluster 7. Ten genes with lowest p_val_adj were presented.

## Discussion

In this study, we performed a time-course single-cell transcriptome profiling of hESC-derived LSCs, and revealed the gene expression patterns and LSCs developmental trajectory. Previous studies showed that *bona fide* LSCs have the potential to establish and maintain long-term corneal repair. Many studies have investigated the identity of human LSCs and several candidate LSCs markers have been identified, such as TP63 (well-known as p63) (Pellegrini et al., 2001), KRT14 (Kurpakus et al., 1994), ITGA6 (Hayashi et al., 2008), NTRK1 (Qi et al., 2008), ABCG2 (Budak et al., 2005; de Paiva et al., 2005), KRT15 (Yoshida et al., 2006), ABCB5 (Ksander et al., 2014). Besides, the terminally differentiated markers KRT3 and KRT12 were absent in LSCs (Gonzalez et al., 2018; Schermer et al., 1986). However, according to our single cell expression profiling data, hESC-derived LSCs showed significantly cellular heterogeneity using current protocols. For example, TP63 expressed cells only accounted for 11.21% of cells at Day 14 and 2.46% at Day 21 (Fig 1E). Therefore, our data suggested that the heterogenic subpopulations should be further characterized and the current hESCs-LSCs differentiation methods need to be optimized accordingly.

Until recently, the developmental origin of LSCs remained elusive (Gonzalez et al., 2018), and LSCs could be developmental descendants of the surface ectoderm as well as the periocular mesenchyme. Our scRNA-seq data revealed that EMT program were activated in the cluster of cells with neural crest characteristics at early hESC-LSCs differentiation stage. During organogenesis, epithelial cells can give rise to mesenchymal cells through EMT while the reverse process, mesenchymal–epithelial transition (MET), can similarly generate epithelial cells (Pei et al., 2019), suggesting LSCs could be differentiated from the periocular mesenchyme through MET. However, our pseudotime trajectory analysis showed that induced mesenchymal cells did not generate LSCs under current culture conditions, and whether the periocular mesenchyme could give rise LSCs remain to be confirmed. Meanwhile, we found excessive cell death occurred in cells cultured in the medium beyond 20 days, indicating the medium used need to be improved for LSCs generation. Nevertheless, our pseudotime analysis identified a hESC-LSCs developmental trajectory. During organogenesis, cell cycle modulation is important for cell fate determination (Budirahardja and Gonczy, 2009). According to our trajectory, cell cycle related genes, such as CCNB1, CDC20, MKI67, and TOP2A, showed variable expression across hESC-LSCs developmental pseudotime (Fig 3F).

For long term restoration of visual function caused by LSCD, LSCs based transplantation either through autologous or allogenic grafting of limbal tissue, or cultured and expanded limbal cells have already shown effectiveness in the treatment (Atallah et al., 2016). However, so far, only TP63 positive LSC cells were reported to be associated with therapeutic success (Rama et al., 2010). But TP63 could not be applied to sort pure population of LSCs, and isolation of pure LSCs is still the bottleneck concerning the clinical application of LSCs. Therefore, other molecular markers are needed for successful prospective enrichment of LSC cells capable of long-term corneal restoration (Gonzalez et al., 2018). Identification of specific biomarkers for isolating and characterizing LSCs is crucial for both understanding their basic biology and translating in clinical application (Gonzalez et al., 2018; Sonam et al., 2019). According to our scRNA-seq data, TP63 expressed LSCs showed relative quiescence compared to their progeny, and genes related to cell cycle were significantly upregulated in highly proliferative progeny (TACs), which are in line with previous reports that epithelial stem cells are relatively quiescent and give rise to TACs (Lavker and Sun, 2003). Besides reported markers - TP63 and ITGA6, TFs such as CXXC5, IRF6, SKIL, NR2F2, IRX4 etc., and CD genes such as SDC1, CD9, IGF1R, ALCAM etc., were newly identified as potential markers that highly expressed in hESC-derived LSCs (Fig 4C and 4D). Thus, these data provided valuable sources for characterization of LSCs and optimization of hESC-LSCs differention protocols.

In summary, we studied the time-course changes during hESC-LSC differentiation *in vitro* at single-cell level, and revealed significant transcriptional heterogeneity. Based on current protocol in this study, expression heterogeneity of reported LSC markers were identified in subpopulations of differentiated cells. EMT has been shown to occur during differentiation process, which could possibly result in generation of untargeted cells. Pseudotime trajectory revealed transcriptional changes and signatures of commitment for LSCs and their progeny (TACs) that derived from pluripotent stem cells. Furthermore, some new potential makers for LSCs were identified, which are valuable for future investigation of elucidating identity and developmental origin of human LSCs.

## Data accession

The data that support the findings of this study have been deposited into CNGB Sequence Archive (CNSA) (Guo et al., 2020) of China National GeneBank DataBase (CNGBdb)(F.; et al., 2020) with accession number CNP0001218.

## Author Contributions

Conceptualization, C.S., and X.Z.; Methodology and Investigation, C.S., H.W., Q.M., C.C., J.Y. and X.Z. Writing, C.S., H.L., B.L., and X.Z.; Funding Acquisition, B.L., and X.Z.

## ACKNOWLEDGMENTS

This work was supported by Science, Technology and Innovation Commission of Shenzhen Municipality under grant No. KQJSCX20170322143848413 to X.Z. and Shenzhen Municipal Government of China under grant No. 20170731162715261 to B. L. The funders had no role in study design, data collection and analysis, decision to publish, or preparation of the manuscript. It was also supported by BGI-Shenzhen. We thank members of China National GeneBank for technical support.

