## Supplemental Figures for "Time-course single-cell RNA sequencing reveals transcriptional dynamics and heterogeneity of limbal stem cells derived from human pluripotent stem cells"

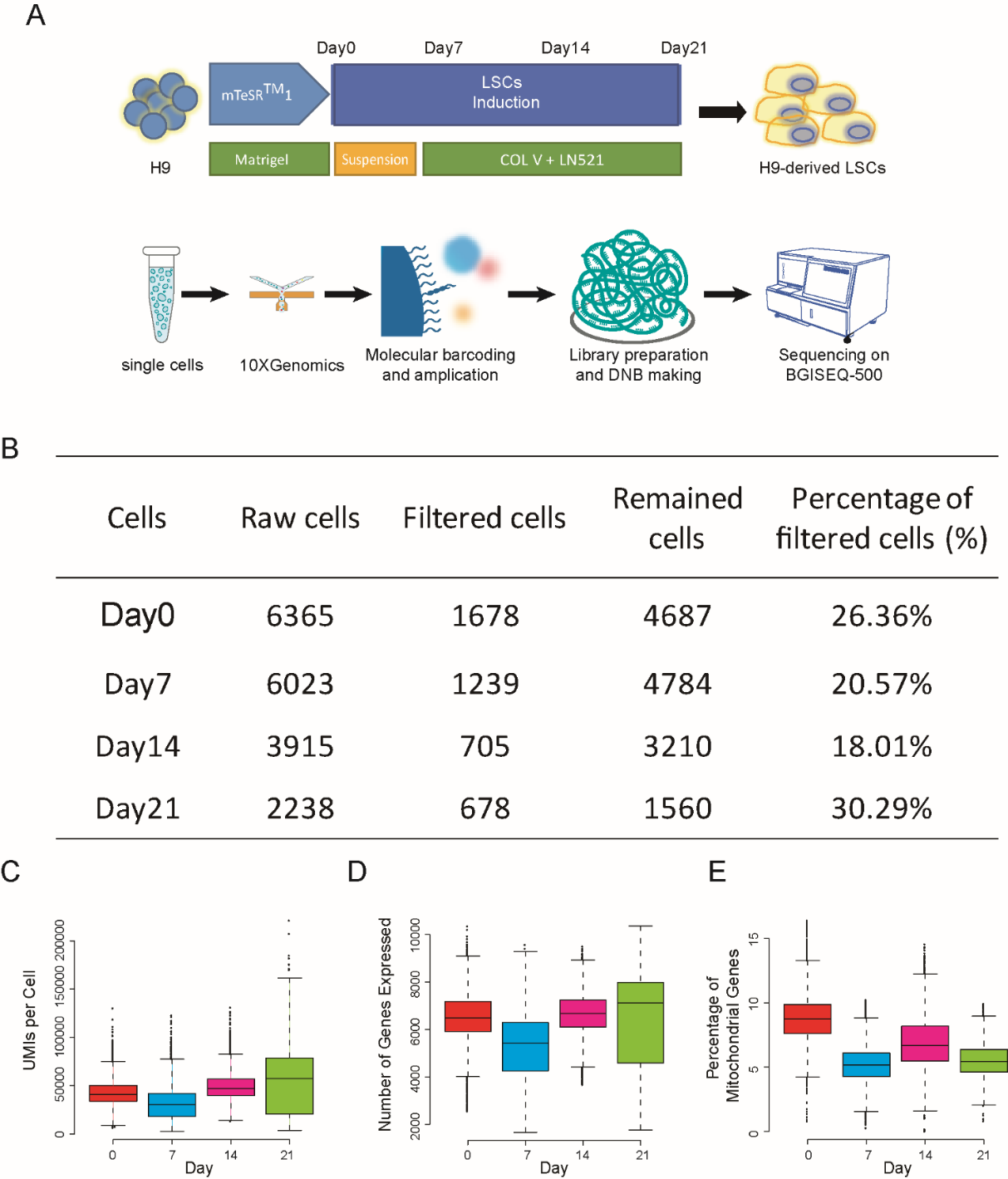

**Fig S1. Overview of the experimental procedure and data quality**

(A) Schematic representation of differentiation procedure and Single cell RNA-seq library preparation and sequencing for hESC-derived LSCs. (B) Number of cells sequenced and filtered for each time. (C) Boxplot showing distribution on UMIs per cell for each time. (D) Boxplot showing distribution on number of genes obtained per cell for each time. (E) Boxplot showing distribution on percentage of mitochondrial genes per cell for each time.

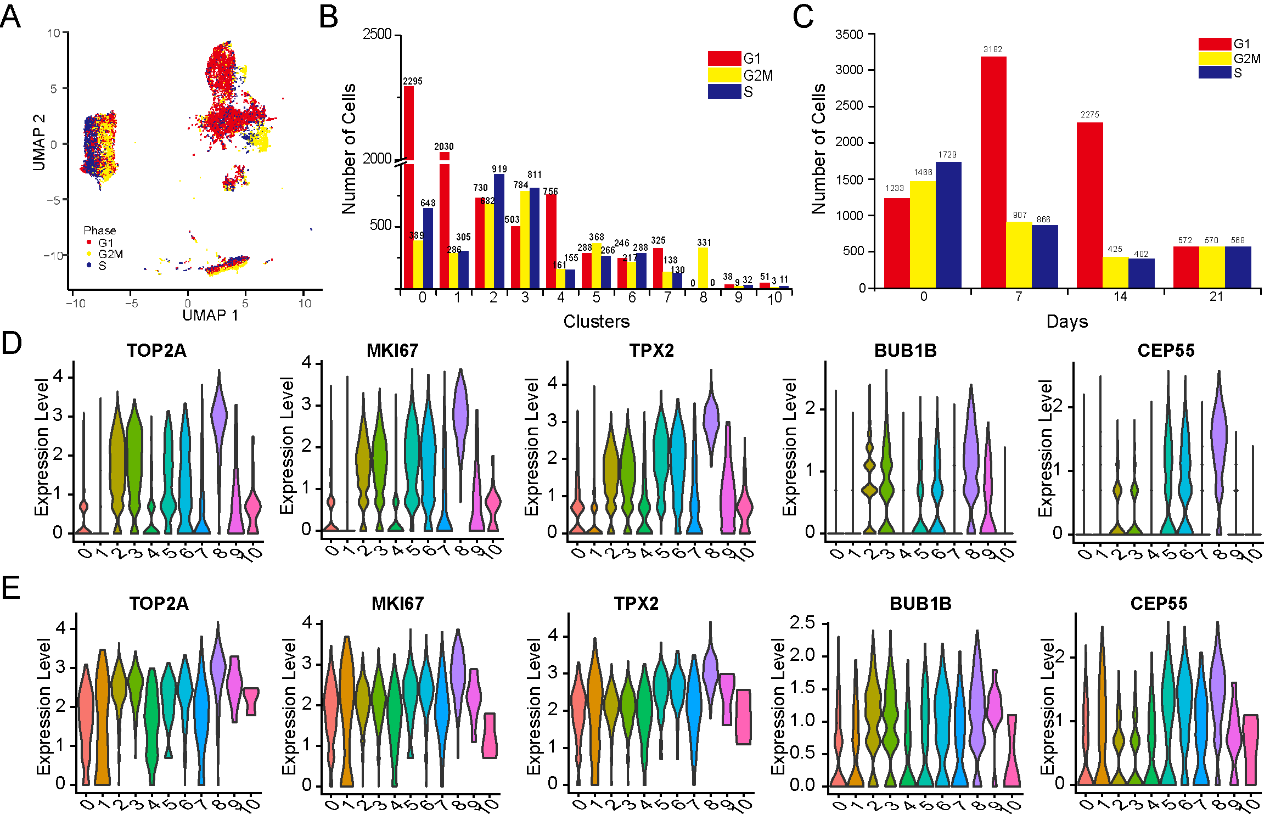

**Figure S2. Pseudotime analysis characterizes expression changes throughout hESCs-derived LSCs differentiation**

(A) UMAP visualizing the results of cell cycle phases assigned for cells sequenced at the four times. (B) Barplot showing number of cells assigned to the cell cycle phases for each cluster. (C) Barplot showing number of cells assigned to the cell cycle phases for each day. (D and E) Violin plot representing expression of cell cycle related genes for only G2M phase cells (D) and all cells (E) in each cluster.

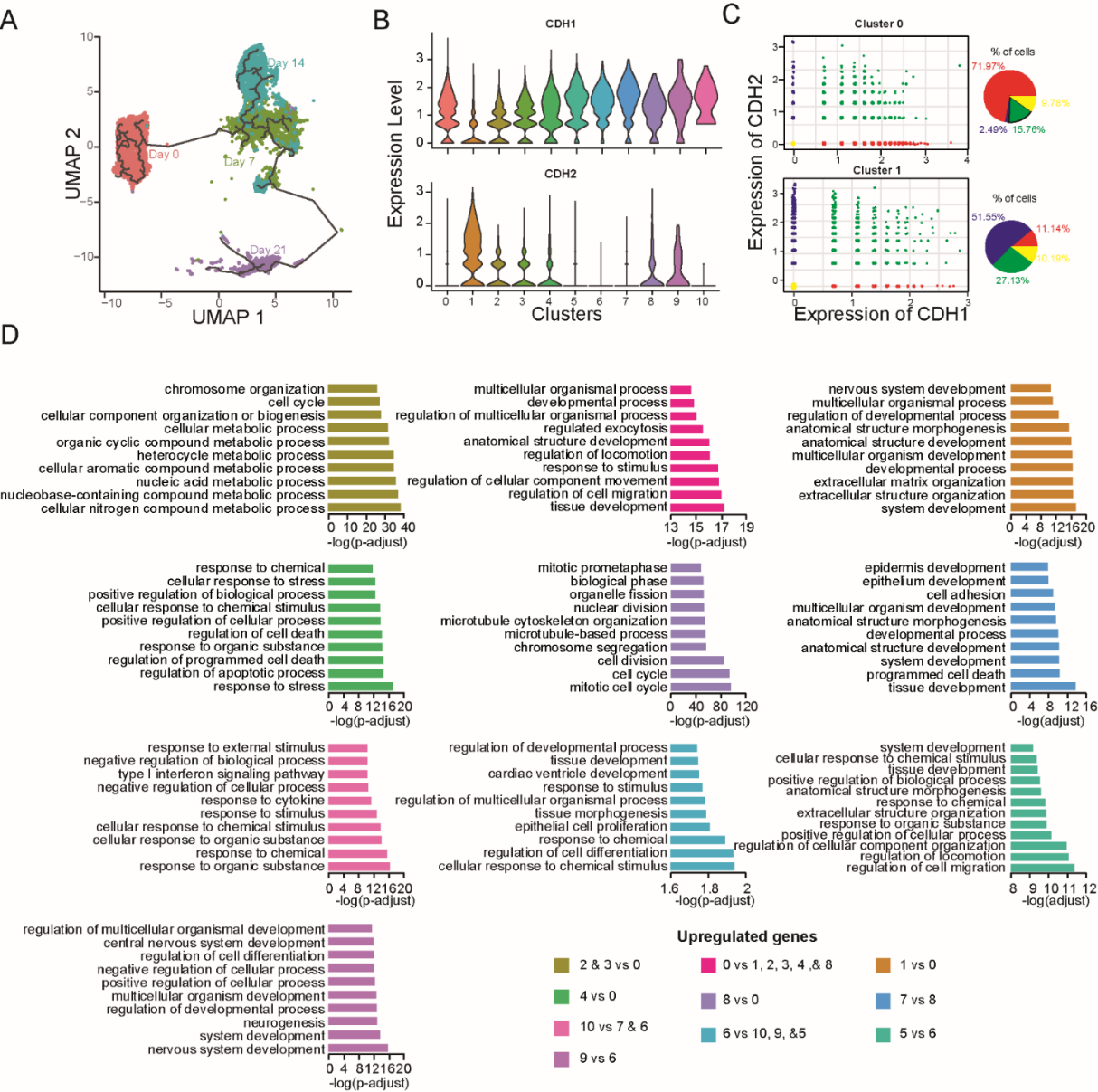

**Figure S3. Pseudotime analysis characterizes expression changes throughout hESCs-derived LSCs differentiation**

(A) UMAP visualizing developmental trajectory of cells in each day. (B) Violin plot showing distribution of expression for CDH1 and CDH2 genes in each cluster. (C) Expression of CDH1 and CDH2 genes in cluster 0 and cluster 1. Scatter plots showing expression of CDH1 and CDH2 in each cell (left) and Pie charts showing percentage cells with different expression pattern of CDH1 and CDH2 genes for cluster 0 and cluster 1. (F) Barplots showing GO biological process enrichment for upregulated genes among adjacent cluster over the trajectory. Only 10 terms with lowest p-adjust values were presented.

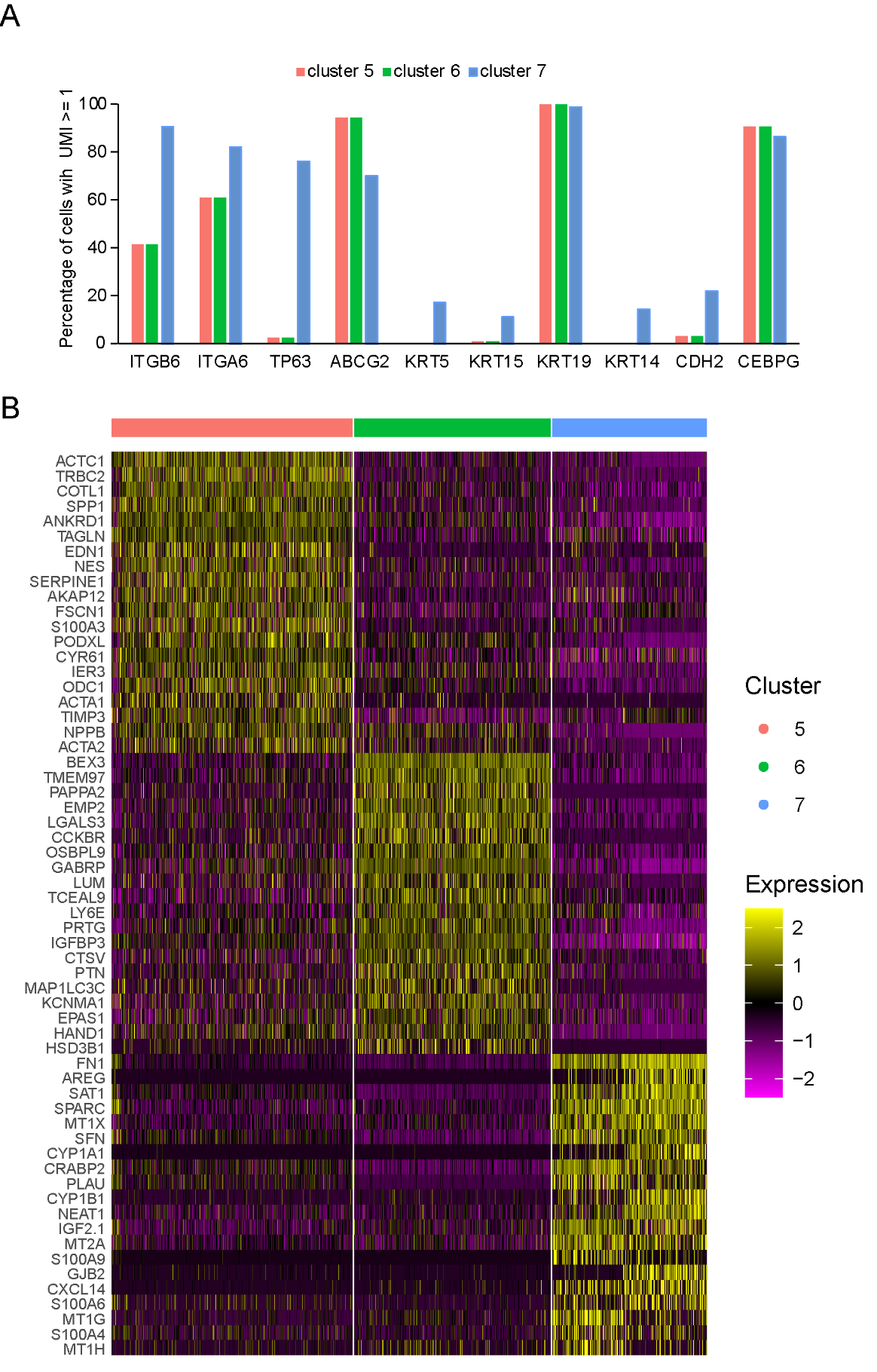

**Figure S4. Transcriptional difference of subpopulations in hESCs-derived LSCs**

(A) Barplot showing percentage of cells expressing (UMI ≥1) some candidate LSCs markers in cluster 5, cluster 6, and cluster 7. (D) Heatmap representing differentially expressed genes among cluster 5, cluster 6, and cluster 7. Twenty genes with lowest p_val_adj were presented.
